# Liposomal encapsulation of polysaccharides (LEPS) as an effective vaccine strategy to protect aged hosts against *S. pneumoniae* infection

**DOI:** 10.1101/2021.10.18.464850

**Authors:** Manmeet Bhalla, Roozbeh Nayerhoda, Essi Y. I. Tchalla, Alexsandra Abamonte, Dongwon Park, Shaunna R. Simmons, Blaine A. Pfeifer, Elsa N. Bou Ghanem

**Affiliations:** Department of Microbiology and Immunology, University at Buffalo, The State University of New York, Buffalo, NY; Department of Biomedical Engineering, University at Buffalo, The State University of New York, Buffalo, NY; Department of Chemical and Biological Engineering, University at Buffalo, The State University of New York, Buffalo, NY; Gene and Tissue Engineering Center, University at Buffalo, The State University of New York, Buffalo, NY

**Keywords:** pneumococcal disease, aging, vaccines, liposomes, immunosenescence, antibodies

## Abstract

Despite the availability of licensed vaccines, pneumococcal disease caused by the bacteria *Streptococcus pneumoniae* (pneumococcus), remains a serious infectious disease threat globally. Disease manifestations include pneumonia, bacteremia, and meningitis, resulting in over a million deaths annually. Pneumococcal disease disproportionally impacts elderly individuals ≥65 years old. Interventions are complicated through a combination of complex disease progression and 100 different bacterial capsular polysaccharide serotypes. This has made it challenging to develop a broad vaccine against *S. pneumoniae*, with current options utilizing capsular polysaccharides as the primary antigenic content. However, current vaccines are substantially less effective in protecting the elderly. We previously developed a Liposomal Encapsulation of Polysaccharides (LEPS) vaccine platform, designed around limitations of current pneumococcal vaccines, that allowed the non-covalent coupling of polysaccharide and protein antigen content and protected young hosts against pneumococcal infection in murine models. In this study, we modified the formulation to make it more economical and tested the novel LEPS vaccine in aged hosts. We found that in young mice (2-3 months), LEPS elicited comparable responses to the pneumococcal conjugate vaccine Prevnar-13. Further, LEPS immunization of old mice (20-22 months) induced comparable antibody levels and improved antibody function compared to Prevnar-13. Importantly, LEPS protected old mice against both invasive and lung localized pneumococcal infections. In summary, LEPS is an alternative and effective vaccine strategy that protects aged hosts against different manifestations of pneumococcal disease.

## 1. Introduction

*Streptococcus pneumoniae* (pneumococcus) is an opportunistic pathogen that asymptomatically resides in the upper respiratory tract of humans but can cause serious life-threatening infections (1, 2). These include pneumonia which can progress to invasive pneumococcal disease leading to bacteremia, meningitis, and endocarditis (3, 4). Pneumococcal infections result in more than a million deaths annually worldwide (1), and in the USA alone, the estimated annual direct medical cost associated with pneumococcal disease is approximately $3.5 billion (5, 6). Pneumococcal infections are more problematic in elderly individuals (above 65 years of age) in terms of both health burden and treatment cost (6, 7). Despite the availability of vaccines and antibiotics, *S. pneumoniae* remains the leading cause of community acquired bacterial pneumonia and nursing home associated pneumonia in the elderly (8-10). Older individuals account for 71% of all pneumococcal cases and 82% of associated deaths (11). The yearly cost associated with hospitalization in this population will increase significantly as the number of elderly individuals is projected to double in the coming decades (12). To further complicate the current health concerns, elderly individuals are more at risk of acquiring drug-resistant infections (13) which is on a rise in *S. pneumoniae* (14).

There are 100 different *S. pneumoniae* serotypes identified based upon capsular polysaccharide content (15, 16), which in part regulates the severity of pneumococcal disease (17). Currently licensed vaccine formulations consist of capsular polysaccharides from the most common disease-causing *S. pneumoniae* serotypes prevalent globally. The pneumococcal polysaccharide vaccine (Pneumovax 23/PPSV23) consists of polysaccharides from 23 pneumococcal serotypes and elicits a T cell-independent antibody response, while the pneumococcal conjugate vaccine (Prevnar 13/PCV13) is comprised of 13 serotypes covalently linked to a diphtheria toxoid CRM197 carrier protein which produces a T cell-dependent immune response in the host (18). There are several limitations to the current vaccines, one of which is the phenomenon of serotype replacement wherein the targeting of a certain and limited number of pneumococcal serotypes by these vaccines has led to an increased prevalence of non-vaccine serotypes (19, 20). In addition, having only polysaccharides-centric vaccine formulations results in serotype-specific antibodies which mainly target nasopharyngeal colonization or the invasive stage of pneumococcal infection during which *S. pneumoniae* upregulate capsule expression to survive the host immune response. This strategy provides only limited host protection given the fact that pneumococcus undergoes transcriptomic changes to alter expression of several factors, including capsule expression, when transitioning from the stage of colonization to pulmonary or systemic infection (21, 22). Moreover, the recent emergence of disease-associated non-encapsulated pneumococcal strains carrying antibiotic resistance genes (23) highlights the necessity of novel vaccine formulations, which in addition to serotype-specific polysaccharide content, also need to include other pneumococcal antigen(s), ideally the ones shared by multiple *S. pneumoniae* serotypes to broaden vaccine coverage.

In response, we have previously developed an alternative vaccine platform termed Liposomal Encapsulation of Polysaccharides (LEPS) which is designed to address the limitations of current vaccine options (24). With LEPS, pneumococcal capsular polysaccharides are localized internally within the liposomal carrier. To date, we have included final LEPS formulations with up to 24 different serotype polysaccharides (25). To mimic conjugated vaccine options (such as PCV13), the LEPS vehicle is engineered with a noncovalent surface attachment mechanism to localize protein components. In the past, we have surface localized both CRM197 and a *S. pneumoniae* protein antigen (PncO) that is upregulated during transition of pneumococci from a benign colonizer to a more invasive pathogen and that is well-conserved across pneumococcal strains (24, 25). The LEPS platform thus allows for a broad degree of serotype coverage, provokes the same immune response provided by conjugate vaccines, and can be tailored to include a protein component that can account for the different stages of pneumococcal disease progression.

The efficacy of currently licensed pneumococcal vaccines, defined as prevention of infection by vaccine serotypes, is limited in the elderly. Although protective against bacteremia, PCV13 and PPSV23 show only 45% and 33% protection, respectively, against pneumonia in older individuals (18). This is largely driven by an overall decline in the immune system with aging, also known as immunosenescence, which results in reduced antibody levels and function following vaccination with PPSV23 (26) and PCV13 (27) in elderly individuals. With the success demonstrated by the LEPS vaccine in young mice, as assessed for serotype coverage and directed protection from *S. pneumoniae* challenge, we conducted the current study to test the LEPS vaccine in aged hosts in preclinical murine models of infection. We present data with old mice that support the prospect of the LEPS platform for effective prevention of pneumococcal disease within aged hosts.

## 2. Materials and Methods

### 2.1 Ethics statement

All animal studies were performed in accordance with the recommendations in the Guide for the Care and Use of Laboratory Animals. Procedures were reviewed and approved by the University at Buffalo Institutional Animal Care and Use Committee.

### 2.2 Mice

All the animal work was done in C57BL/6 young (2-3 months) and old (22-24 months) male mice. The mice were purchased from Jackson Laboratories (Bar Harbor, Maine) and the National Institute on Aging colonies and housed in a specific-pathogen-free facility at the University at Buffalo.

### 2.3 Bacterial strains and growth conditions

*Streptococcus pneumoniae* serotypes 4 (TIGR4 strain) and 19F (P1084 strain) were a kind gift from Andrew Camilli (Tufts University). Bacteria were grown to mid-log phase (corresponding to an OD_650nm_ of 0.7-0.8) in Todd–Hewitt broth (BD Biosciences) supplemented with Oxyrase and 0.5% yeast extract at 37°C/5% carbon dioxide. Aliquots were frozen at –80°C in growth media with 20% glycerol. Prior to use, aliquots were thawed on ice, washed and diluted in phosphate buffered saline to required numbers. Bacterial titers were confirmed by plating on tryptic soy agar plates supplemented with 5% sheep blood agar (Hardy Diagnostics).

### 2.4 Protein production and purification

Recombinant production of the PncO protein antigen was accomplished as reported previously (24, 25, 28, 29). Briefly, *Escherichia coli* strain BL21(DE3) containing a plasmid with the PncO gene was grown in lysogeny broth (LB) medium with ampicillin (100 μM) while shaking overnight at 37°C. The overnight culture was diluted 1:1000 into LB (with ampicillin) and grown under the same conditions to an OD_600nm_ of 0.4-0.5. The culture was then induced by adding isopropyl β-D-1-thiogalactopyranoside (IPTG; 300 μM) and incubated with shaking at 30°C for six hours. The cell culture was harvested by centrifugation for 15 min at 3,200 rcf at 4°C. The resulting pellet was resuspended gently in Buffer A (50 mM Na_2_HPO_4_, 500 mM NaCl, and 10% glycerol (pH 7.5), placed in ice, and the suspended cells lysed by sonication using a small, tapered tip in 3 cycles of 45 s on and 60 s off at 50 Amp, 20 kHz. The post-sonication solution was centrifuged for 15 min at 12,000 rcf at 4°C. The supernatant was maintained at 4°C for protein purification with a prepared HisTrap HP column (GE Healthcare). Purified protein samples were concentrated using an Amicon 4 centrifugal filtration tube centrifuged for 5 min at 3600 rcf (4 °C), and the final protein concentration was measured via Bradford analysis (Thermo Fisher).

### 2.5 LEPS formulation, assembly, and assessment

LEPS formulation and assessment was completed as previously reported (24, 25, 28, 29). Briefly, all lipids were purchased from Avanti, Thermo Fisher, or Sigma Aldrich, and pneumococcal polysaccharides serotype 19F and 4 were obtained from ATCC. 1:2-dioleoyl-sn-glycero-3-phospho-(1’-rac-glycerol) (DOPG), 1,2-dioleoyl-sn-glycero-3-phosphocholine (DOPC), 1,2-dioleoyl-sn-glycero-3-[(N-(5-amino-1-carboxypentyl)iminodiacetic acid)succinyl] (nickel salt) (DGS-NTA(Ni)), 1,2-distearoyl-sn-glycero-3-phosphoethanolamine-N-[amino(polyethylene glycol)-2000] (ammonium salt) (DSPE-PEG(2000)), and cholesterol were dissolved in chloroform with the molar ratio of 3:3:1:0.1:4. Then, 19F and 4 solutions with concentrations of 2.2 μg per dose were separately added to different lipid mixtures and vortexed for 1 minute. Both lipid mixtures were fully evaporated using a rotatory evaporator until a thin film was formed and then rehydrated with PBS at 45°C to dissolve the film prior to extrusion through a 200 nm pore size membrane of a handheld extruder. The resulting liposomes encapsulating the polysaccharides were separated from free floating and unencapsulated polysaccharides by centrifugation for 5 minutes at 1,200 rcf and 4°C using a centrifugal tube with a 300 kDa (Pall Co) filtration device. After readjusting to the initial volume with PBS, 34 μg per of the PncO protein was incubated with the liposome solution for 30 minutes at room temperature. The centrifugation step with filtered tubes was repeated twice to separate the unbound protein, resulting in the final formulation of liposome encapsulating PS with bound protein on the liposomal surface. To this final formulation, 125 μg per dose of aluminum phosphate was added.

### 2.6 Mice immunization and sera collection

Mice were immunized via intramuscular injection into the caudal thigh muscle (30) with 50 μL of the following vaccine formulations: LEPS containing PncO, alum and either serotype 4 or 19F capsular polysaccharide antigen (LEPS); Prevnar 13 (PCV13) (Wyeth Pharmaceuticals); or the controls PBS (Sham) and empty LEPS vector (Empty LEPS plus alum). PCV13 was administered as a single dose, as approved for human use. Booster doses of LEPS or Empty LEPS were given at two weeks following the first inoculation. Sera from each mouse were collected from the tail vein at weeks 2 and 4 following the initial vaccination and saved at –80°C for quantification of antibody titers (31). Sera from each mouse were collected via cardiac puncture 4 weeks following the initial vaccination, pooled per group and saved at –80°C for quantification of antibody function.

### 2.7 Measurement of antibody titer

Enzyme-linked immunosorbent assay (ELISA) was used to determine the antibody titer from the sera samples, as previously described (30, 32). Briefly, Nunc maxisorp plates were coated overnight at 4°C with type 4 pneumococcal polysaccharide (ATCC) or type 19F pneumococcal polysaccharide (ATCC), each at 2 μg/well, or heat killed *S. pneumoniae* TIGR4 strain at 2 × 10^5^ colony forming units (CFU)/well. For type 4 and type 19F polysaccharide conditions, the sera were preabsorbed with a pneumococcal cell wall polysaccharide mixture (CWP-multi, Cederlane) to neutralize noncapsular Abs and added to the plate. Following incubation with sera samples, pneumococcal-specific Abs were detected using horseradish peroxidase–conjugated goat anti-mouse immunoglobulin M (IgM; Invitrogen), IgG (Millipore Sigma), or IgG1, IgG2b, IgG2c, or IgG3 (Southern Biotech) followed by addition of TMB substrate (Thermo Scientific). Readings were taken at OD_650_ using a BioTek reader with a program set for kinetic ELISAs where readings were taken every minute for a total of 10 minutes. Antibody units were calculated as percentages of a hyperimmune standard sera included in each ELISA. Hyperimmune standard sera were generated as previously described (31, 32). Briefly, C57BL/6 young mice were intranasally inoculated with *S. pneumoniae* TIGR4 or 19F over the period of 4 weeks with once weekly inoculations, as previously described (32). Mice were then immunized with PCV13 at week 4. Approximately, 4 weeks following PCV13 vaccination, mice were euthanized, sera collected and pooled, and aliquots stored at −80°C. Total antibody levels in the circulation were then determined in sera from sham treated controls (naïve) for each age group and the hyperimmune standard serum using antibody quantification kits from Invitrogen for IgG (88-50400), IgM (88-50470), IgG1 (88-50410), IgG2b (88-50430), IgG2c (88-50670), and IgG3 (88-50440). To calculate the total amounts of anti-pneumococcal antibodies, the antibody levels in the naïve sera was subtracted from the hyperimmune standard sera and total antibody levels in the different sera samples were then extrapolated using the measured percentages of the hyperimmune standard sera.

### 2.8 PMN isolation

PMNs were isolated from the bone marrow of naïve young and old C57BL/6 mice through density gradient centrifugation, using Histopaque 1119 and Histopaque 1077, as previously described (31, 33). Following isolation, the PMNs were resuspended in Hanks’ Balanced Salt Solution (HBSS)/0.1% gelatin without Ca^2+^ and Mg^2+^ and kept on ice until used in subsequent assays.

### 2.9 Opsonophagocytic killing

The opsonic capacity of antibodies in the sera was determined using an opsonophagocytic (OPH) killing assay with primary PMNs, as previously described (31, 33-37). Briefly, 1 × 10^3^ bacteria (*S. pneumoniae* TIGR4) grown to mid-log phase were incubated with 3% sera from mice immunized with the LEPS formulation containing serotype 4 capsular polysaccharide antigen, PCV13, or the controls (Sham or Empty LEPS vector). Reactions were rotated for 40 min at 37°C to allow opsonization. Following opsonization, bacteria were mixed with 1 × 10^5^ PMNs in 100 μL reactions of HBSS/0.1% gelatin. Reactions were then rotated for 45 min at 37°C. Following incubation, the plates were kept on ice for 2 min to stop the reaction. The ability of sera from each treatment group to induce opsonophagocytic killing of bacteria by PMNs was then determined by plating the reaction mixtures on blood agar plates and comparing colony counts and calculating the percent of bacteria killed with respect to a no PMN control under the exact sera conditions.

### 2.10 Binding assay using H292 cells

The ability of sera to neutralize the binding of *S. pneumoniae* to pulmonary epithelium was determined through an assay using human pulmonary mucoepidermoid carcinoma-derived NCI-H292 (H292) cells (ATCC), as previously described (38). H292 cells were grown and maintained following a previously described protocol (38, 39). Approximately, 2.5 × 10^5^ epithelial cells were seeded in tissue culture-treated flat bottom 96-well plates (Corning) and allowed to adhere overnight. The following day, cells were washed three times with PBS and infected with the *S. pneumoniae* TIGR4 strain at a multiplicity of infection (MOI) of 10 in antibiotic-free media. Prior to infection, bacteria were opsonized for 40 min at 37°C with 10% sera from mice immunized with serotype 4 LEPS, PCV13, or the controls (Sham or Empty LEPS). The reaction plates were spun down to facilitate cell-bacterial interaction and incubated for 1 hr at 37°C/CO_2_. The cells were then washed five times with PBS to remove unbound bacteria, lifted with 0.05% trypsin/ethylenediaminetetraacetic acid (EDTA) (Invitrogen), and mixed vigorously to produce a homogeneous solution. Serial dilutions were then plated on blood agar plates to determine the bacterial CFU. The percent of bacteria bound was determined with respect to a no cell control of the same sera condition where bacteria were added to the wells and incubated for an hour under the same experimental conditions. The number of bacteria bound to H292 cells in the Sham sera condition was set as 100% and relative changes in bacterial binding were then calculated for other sera conditions.

### 2.11 Animal infections

Mice were infected with *S. pneumoniae*, as previously described (32, 35). Briefly, mice were intratracheally (i.t) challenged with either the serotype 4 (5 × 10^5^ CFU) or 19F (2 × 10^7^ CFU) strain of *S. pneumoniae*. Following infection with the *S. pneumoniae* serotype 4 strain, mice were monitored daily over 2 weeks for survival and clinical signs of disease (weight loss, activity level, posture, and breathing, scored healthy [0] to severely sick [21], as previously described (32, 35)) while bacteremia was determined for up to 72 hr post infection. In case of infection with the *S. pneumoniae* serotype 19F strain, mice were scored for clinical signs of disease at twenty-four hours post infection and euthanized to assess bacterial burden in the lung. Lung homogenates and blood were assessed for CFU by plating on blood agar plates.

### 2.12 Statistical analysis

All statistical analysis was performed using Prism9 (Graph Pad). Data with bacterial numbers in blood and lungs were log-transformed to normalize distribution. Bar graphs represent the mean values +/-standard deviation (SD) and line graphs represent the mean values +/-95% confidence interval of the mean (CI). Significant differences were determined with one-way ANOVA followed by Tukey’s multiple comparisons test as indicated. Survival was analyzed using the log-rank (Mantel-Cox) test. Differences between fractions were determined by Fisher’s exact test. All *p* values < 0.05 were considered significant.

## 3. Results

### 3.1 LEPS and Prevnar-13 vaccination induce comparable antibody production in young hosts against *S. pneumoniae*

We had previously shown that a LEPS formulation constructed with surface localized CRM197, to mimic conjugated vaccine options, conferred a similar degree of immunogenicity and protection against *S. pneumoniae* as the pneumococcal conjugate vaccine Prevnar-13 (PCV13) in young mice (24). In an effort to simplify and economize, we altered the LEPS vaccine by replacing CRM197 with the pneumococcal protein antigen PncO (Fig. 1A). To test if the modified LEPS vaccine is immunogenic and still mimics the immunological outcome of PCV13, we first examined the antibody production in young (2 months old) C57BL/6 mice. Mice were immunized by intramuscular injection with the following vaccine formulations (i) LEPS containing capsular polysaccharide conjugated with PncO with alum as adjuvant (LEPS); (ii) PCV13; (iii) a PBS control (Sham); and (iv) an empty LEPS control with alum (Empty LEPS), following the schedule presented in Fig. 1B. Sera from each mouse was collected over time to measure antibody levels (timeline in Fig. 1B).

**Figure 1.**
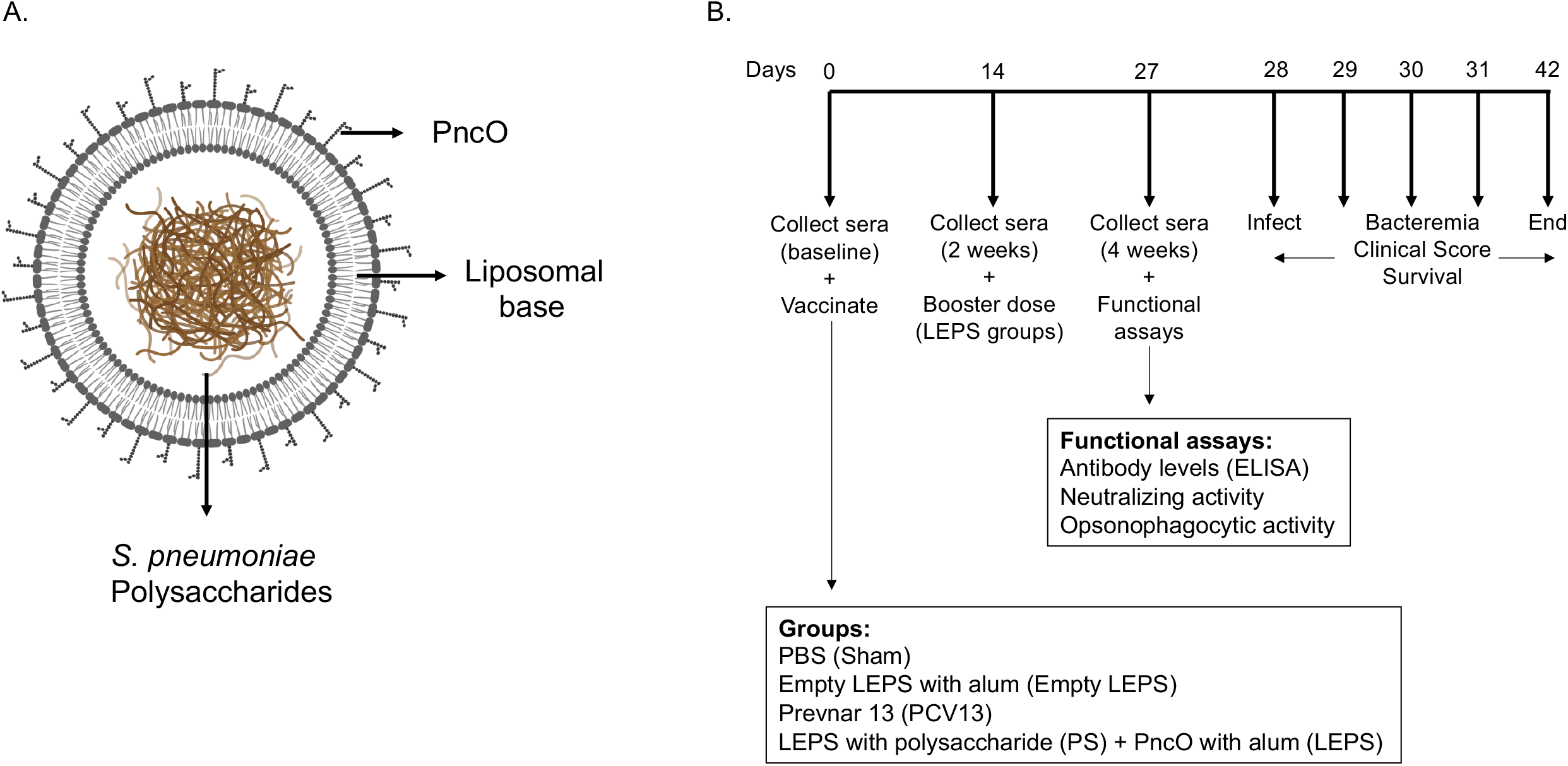
Vaccine design and experimental set-up. (A) LEPS design. The LEPS vaccine consists of the liposomal carrier encapsulating bacterial polysaccharide with PnCO linked to the surface. (B) Vaccination timeline. Mice were mock treated with 50 μL PBS (Sham), empty LEPS with alum (Empty LEPS), or administered either the pneumococcal conjugate vaccine Prevnar 13 (PCV13) or the LEPS vaccine containing bacterial polysaccharide in addition to PnCO and alum (LEPS) via intramuscular (*i*.*m*) injections to the hind legs. Two weeks following the initial vaccination, the LEPS and Empty LEPS groups received a booster dose. Sera were collected at baseline and two and four weeks with respect to the initial vaccination to assess antibody levels and function. Four weeks following vaccination mice were challenged intra-tracheally (*i*.*t*) with *S. pneumoniae* and monitored for bacterial burden and clinical signs of disease.

Serum antibody response was measured using ELISAs against purified polysaccharide. We found that LEPS vaccination was able to induce IgM production at levels that were significantly higher than the Empty LEPS and that were comparable to those induced by PCV13 (Fig. 2A). An important feature of the PCV13 vaccine is the presence of the covalently attached CRM197 protein, which induces T-cell-mediated IgG class switching. To confirm that the LEPS vaccine still elicited IgG class switching (with only PncO as the noncovalently affixed protein conjugate), we next measured total antibody levels as well as the different subtypes of IgG produced. We found that while no IgG was detected in the Empty LEPS treated controls (Fig. 2B), LEPS vaccination induced IgG production that was comparable to PCV13 (Fig. 2B). Similar IgG and IgM responses were also detected against heat-killed bacteria (Fig. S1). When we compared IgG subtypes, we found that LEPS and PCV13 triggered similar class switching to predominantly the IgG3 and IgG2b subtypes (Fig. 2C-F). Overall, these findings confirm that the LEPS vaccine is immunogenic in young hosts and elicits antibody production comparable to PCV13.

**Figure 2.**
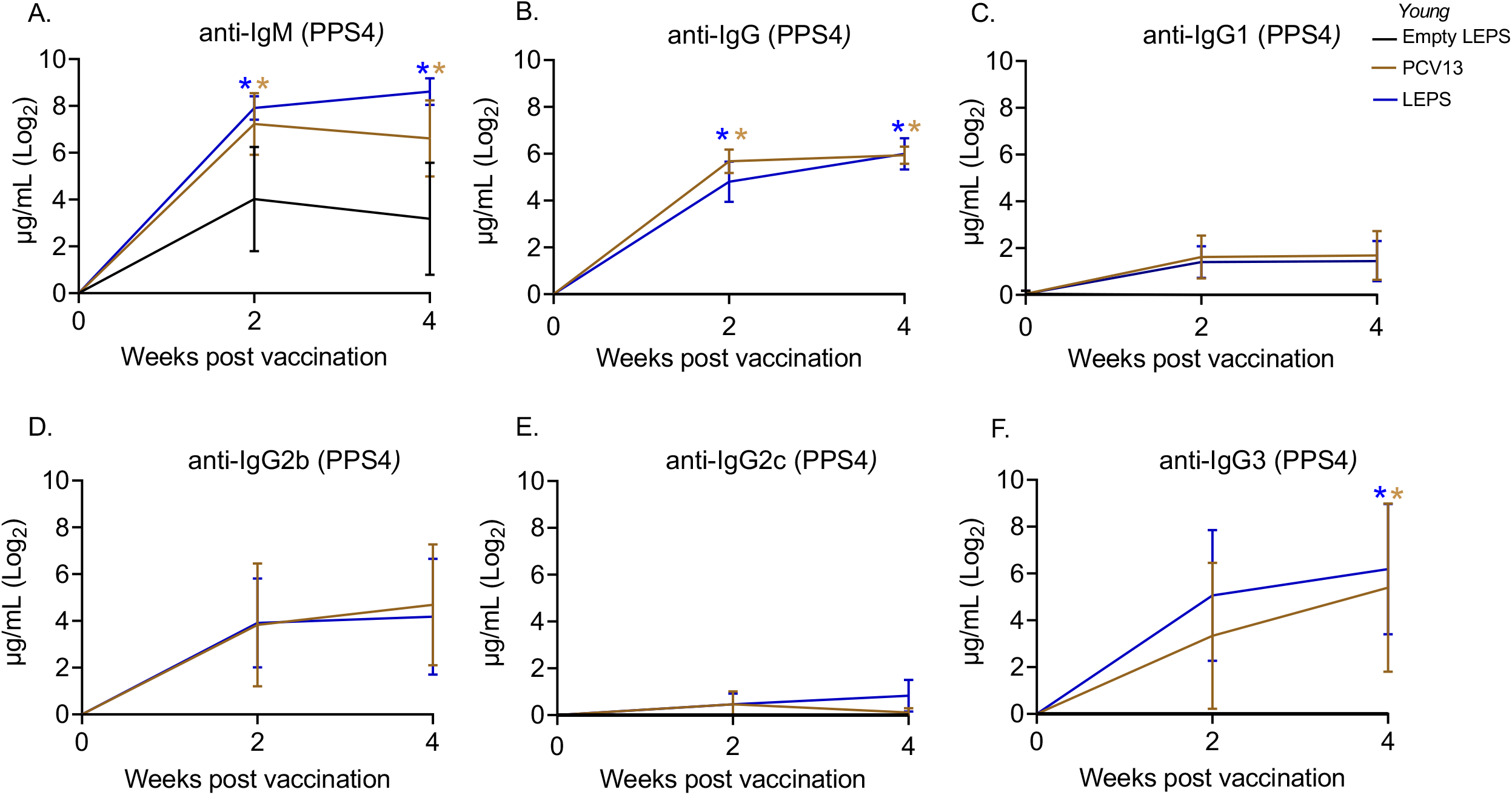
The LEPS vaccine and Prevnar-13 induce comparable levels of antibodies against pneumococcal polysaccharides in young hosts. Young C57BL/6 mice (2-months) were injected *i*.*m*. with LEPS containing serotype 4 capsular polysaccharide antigen, PCV13, or the controls (Sham or Empty LEPS vector) following the timeline in Fig. 1B. Total levels of IgM (A), IgG (B), and IgG subtypes including IgG1 (C), IgG2b (D), IgG2c (E) and IgG3 (F) against purified polysaccharide serotype 4 in the sera were measured over time by ELISA. *p* values were determined by one-way ANOVA followed by Tukey’s test. Asterisks (*, *p*<0.05) denote significant differences between the indicated group and the empty LEPS group. Data shown are presented as mean ± CI and are pooled from 3 separate experiments with n= 10 mice for Empty LEPS, n=10 mice for LEPS, and n=15 mice for the PCV13 groups.

### 3.2 The LEPS vaccine is immunogenic in aged hosts

Aging is accompanied by immunosenescence which is known to blunt immune responses to vaccines (40-42). Given our positive results when using young mice, we next tested if the LEPS vaccine could similarly prompt an immunogenic response in aged hosts. Thus, old (18-22 months) C57BL/6 mice were immunized with the same vaccine formulations used in young mice (Fig. 1B) and antibody levels against purified polysaccharide were measured. Similar to what we observed in young mice, we found that LEPS vaccination of old mice elicited IgM and IgG production comparable to that of PCV13 (Fig 3A and B). Further, LEPS and PCV13 also induced class switching to predominantly IgG2b and IgG3 in old mice (Fig 3C-F). Overall, these data indicate that the LEPS vaccination is immunogenic in old mice and the antibody response mounted to LEPS by aged hosts is comparable to that observed in young controls.

**Figure 3.**
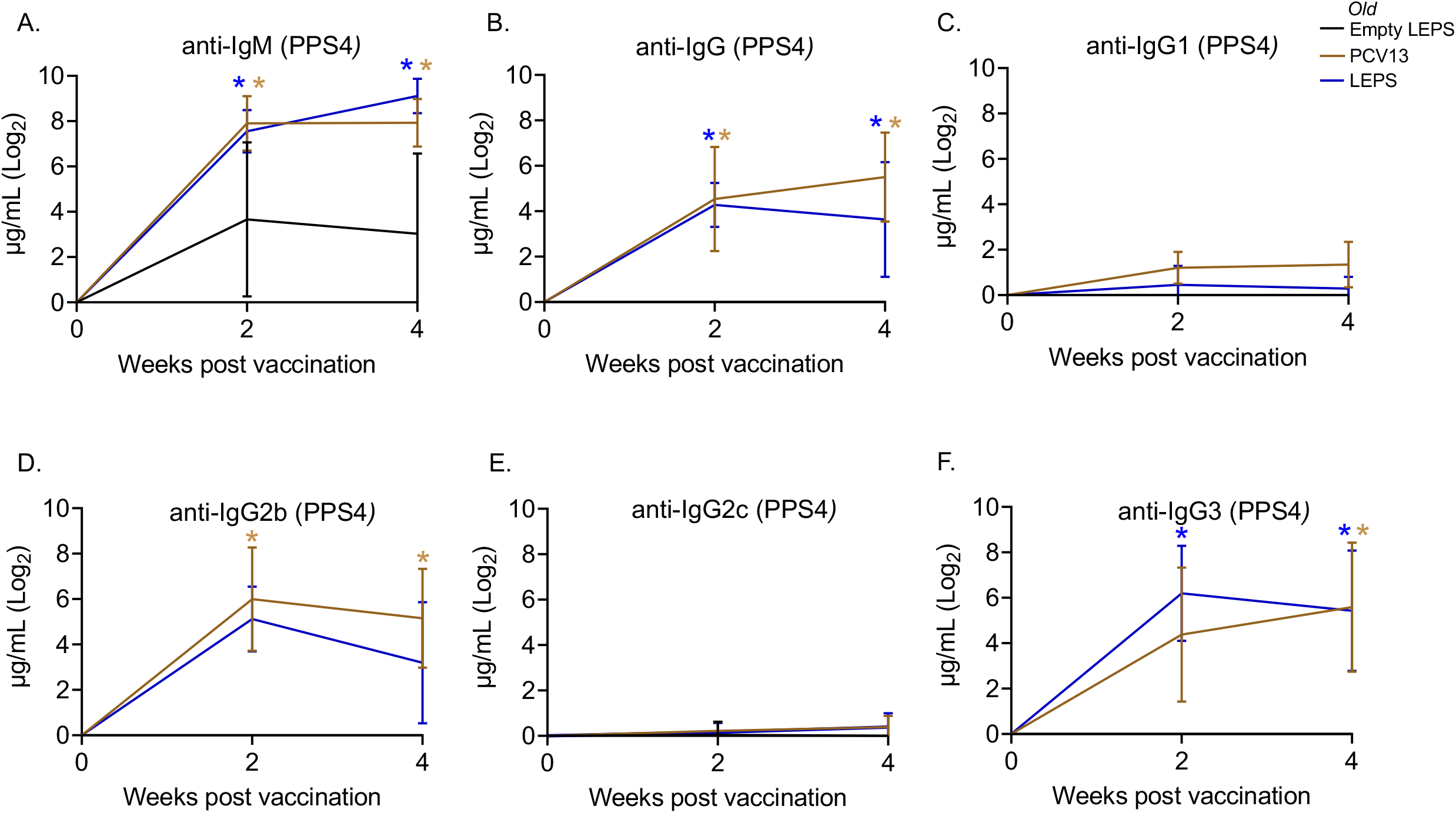
The LEPS vaccine and Prevnar-13 induce comparable levels of antibodies against pneumococcal polysaccharide in old hosts. Old C57BL/6 mice (18-22 months) were injected *i*.*m*. with LEPS containing serotype 4 capsular polysaccharide antigen, PCV13, or the controls (Sham or Empty LEPS vector) following the timeline in Fig. 1B. Total levels of IgM (A), IgG (B), and IgG subtypes including IgG1 (C), IgG2b (D), IgG2c (E) and IgG3 (F) against purified polysaccharide serotype 4 in the sera were measured over time by ELISA. *p* values were determined by one-way ANOVA followed by Tukey’s test. Asterisks (*, *p*<0.05) indicate significant differences between the indicated group and the empty LEPS group. Data shown are presented as mean ± CI and are pooled from 3 separate experiments with n= 5 mice for Empty LEPS, n=10 mice for LEPS, and n=10 mice for the PCV13 groups.

### 3.3 LEPS vaccination elicits enhanced serum neutralizing activity against *S. pneumoniae* in aged hosts

Apart from antibody levels, antibody function is crucial for vaccine efficacy (43). We therefore wanted to assess antibody function following LEPS vaccination. Binding of *S. pneumoniae* to the airway epithelium is an important step that is crucial for host colonization and infection (2, 4, 38, 44). As such, we tested the ability of antibodies induced by the LEPS vaccine to neutralize bacterial binding to H292 cells (type I and II pneumocytes), a human lung epithelial cell line extensively used to study host-pathogen interactions (38, 39, 45, 46). To do so, we compared the ability of sera collected four weeks post-vaccination across the different immunization groups (Fig. 1B) to block bacterial binding relative to the Sham control group (Fig. 1B). We found that in young mice, sera from both the PCV13 and LEPS vaccinated groups significantly reduced the binding of a *S. pneumoniae* serotype 4 strain (serotype covered by both vaccines) to H292 cells by 25% (Fig. 4A). Importantly, sera from the LEPS group caused a significant 2-fold reduction in bacterial binding compared to the Empty LEPS control (Fig 4A). The effects observed were not due to the direct bactericidal activity of the sera as all binding is calculated with respect bacteria incubated with sera alone for each opsonization condition (see Materials and Methods). These data suggest that antibodies induced by both PCV13 and LEPS in the young mice are capable of neutralizing bacterial adherence to host cells.

**Figure 4.**
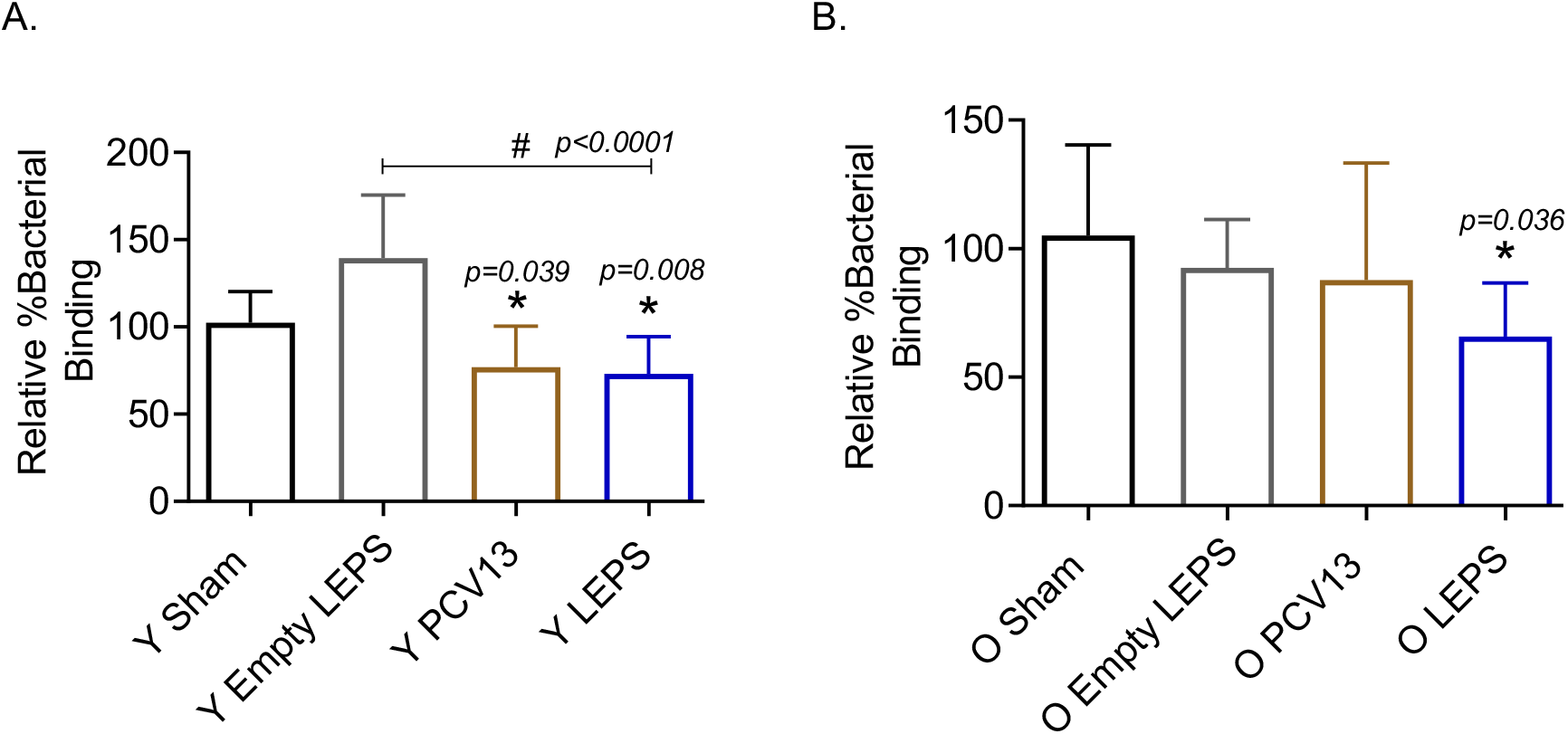
Sera from old mice vaccinated with LEPS, but not Prevnar-13, neutralize the ability of *S. pneumoniae* to bind pulmonary epithelial cells. Sera were collected from mice immunized with LEPS containing serotype 4 capsular polysaccharide antigen, PCV13, or the controls (Sham or Empty LEPS vector) at four weeks post vaccination following the timeline indicated in Fig. 1B. The ability of sera to neutralize bacterial binding to pulmonary epithelial cells was determined. *S. pneumoniae* serotype 4 TIGR4 strain was pre-opsonized for 45 minutes with 10% sera from young (Y) (A) or old (O) (B) mice and used to infect H292 cells for 1 hour at a MOI of 10. The number of bound bacteria was determined by plating on blood agar plates and the percent bacterial binding calculated with respect to bacteria incubated with sera alone for each opsonization condition. The effect of sera on bacterial binding was then determined relative to the Sham group. Asterisks (*) indicate significant differences with respect to the Sham group and hash signs (#) indicate significant differences between the indicated groups as calculated by one-way ANOVA followed by Tukey’s test. Data shown are pooled from four separate experiments where each condition was tested in quadruplicate and presented as mean ± SD.

The function of antibodies is known to decline with age (47). In fact, when we measured the ability of sera isolated from old mice to neutralize bacterial binding, we found that unlike what we observed in young hosts, sera from old mice vaccinated with PCV13 had no effect on pneumococcal binding to H292 cells compared to the sham group (Fig. 4B). Interestingly, when *S. pneumoniae* was opsonized with sera from old LEPS vaccinated mice, there was a significant 40% and 25% reduction in bacterial binding compared to the Sham and Empty LEPS controls, respectively (Fig. 4B). This finding suggests that compared to PCV13, LEPS vaccination induces antibodies that are better at neutralizing bacterial binding to the pulmonary epithelium.

### 3.4 LEPS vaccination elicits enhanced serum opsonic activity against *S. pneumoniae* in aged hosts

The opsonic capacity of antibodies is an important correlate of vaccine protectiveness (48). We therefore measured the ability of sera isolated at four weeks post-immunization across the different groups (Fig. 1B) to induce opsonophagocytic (OPH) killing of a *S. pneumoniae* serotype 4 strain (serotype covered by both vaccines) by primary PMNs from naïve mice, using assays we had previously established (31). We found that in young hosts, compared to the control groups, we saw a significant increase in pneumococcal killing by PMNs when the bacteria were opsonized with sera from PCV13 or the LEPS vaccinated groups, with higher opsonophagocytic killing induced by the latter (Fig 5A). This suggests that the opsonic activity of sera induced by LEPS immunization matched or exceeded that elicited by PCV13.

**Figure 5.**
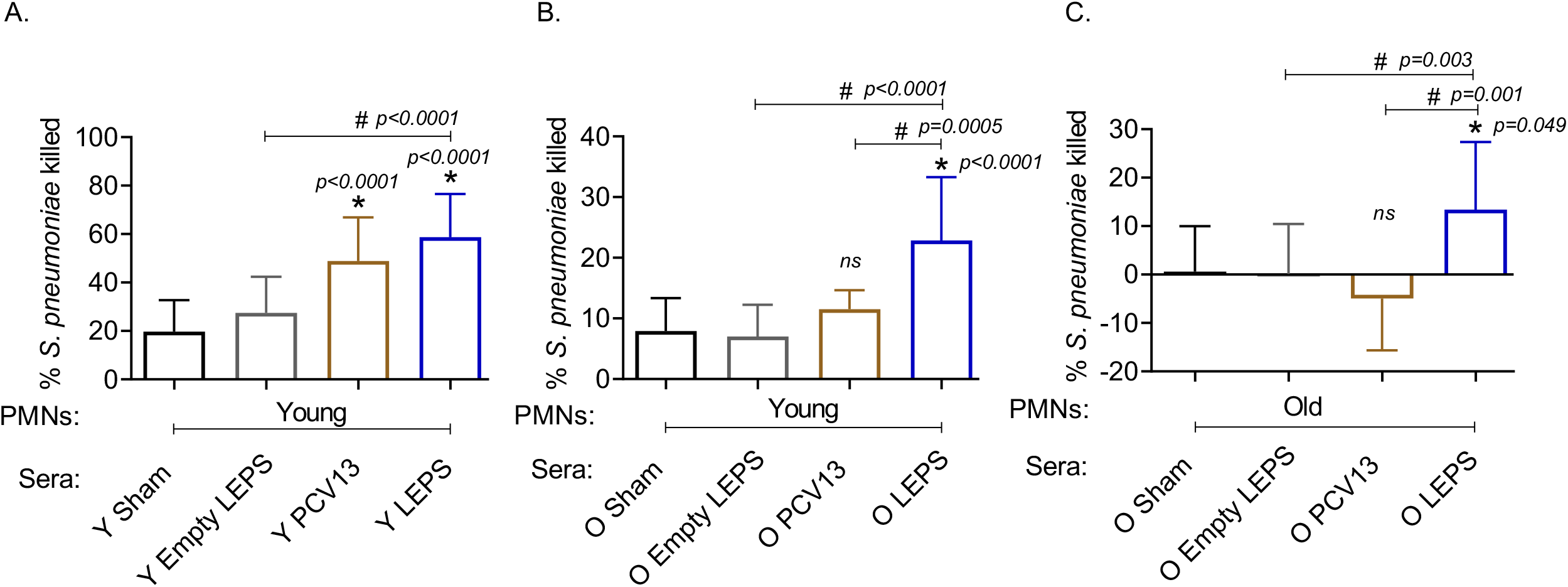
LEPS vaccination of old mice induces sera with better opsonic activity compared to Prevnar-13 vaccination. Sera were collected from young (Y) or old (O) mice immunized with LEPS containing serotype 4 capsular polysaccharide antigen, PCV13, or the controls (Sham or Empty LEPS vector) at four weeks post vaccination following the timeline indicated in Fig. 1B. The ability of sera to induce opsonophagocytic killing of bacteria by PMNs was determined. PMNs were isolated from the bone marrow of naïve young (2 months) (A and B) or old (18-22 month) (C) C57BL/6 mice and mixed for 45 min at 37°C with *S. pneumoniae* serotype 4 TIGR4 strain pre-opsonized with 3% sera from the indicated groups. Reactions were stopped on ice, and viable CFU were determined after serial dilution and plating. The percentage of bacteria killed was determined with respect to a no PMN control for each condition. Asterisks (*) indicate significant differences with respect to the Sham group (ns: not significant) and hash signs (#) indicate significant differences between the indicated groups as calculated by one-way ANOVA followed by Tukey’s test. Data shown are pooled from three separate experiments (n=3 biological replicates or mice per group) where each condition was tested in triplicate (n=3 technical replicates) per experiment and presented as mean ± SD.

We then compared the opsonic activity of sera from old mice. As PMNs function is known to decline with age and can confound the interpretation of the data (35, 49), we first measured the ability of sera isolated from old mice to elicit opsonophagocytic bacterial killing by PMNs isolated from young mice. We found that while sera from PCV13 immunized old mice failed to significantly improve bacterial killing by PMNs relative to Sham controls (Fig 5B), sera from LEPS vaccinated old mice elicited significantly higher opsonophagocytic killing by PMNs than both controls as well as the PCV13 group (Fig 5B). To best mimic *in vivo* conditions, we then age matched the sera and PMNs and measured the ability of sera from old hosts to induce bacterial killing by PMNs from old mice. As expected (35), sera from the control groups (Sham and Empty LEPS) failed to elicit opsonophagocytic bacterial killing by PMNs from old mice (Fig. 3C). Surprisingly, when PMNs were challenged with *S. pneumoniae* opsonized with PCV13 sera from old mice, we failed to see bacterial killing and in fact observed bacterial growth in the presence of PMNs (Fig 5B). In contrast, we saw a significant 10-fold increase in bacterial killing by PMNs in comparison to the control groups when sera from the LEPS vaccinated old mice were used to opsonize *S. pneumoniae* (Fig. 5B). Further, PMN-mediated bacterial killing induced by sera from the LEPS group was significantly higher than that seen with sera from the PCV13 group (Fig 5B). These findings suggest that compared to PCV13, LEPS vaccination induces antibodies that are better at eliciting opsonophagocytic bacterial killing by immune cells.

### 3.5 Young mice immunized with the LEPS vaccine show resistance to invasive pneumococcal infection

We next wanted to test whether the LEPS vaccination confers host protection against pneumococcal infection. *S. pneumoniae* strains can vary (50) and most infections result in primarily pneumonia, but up to 30% of patients with pneumococcal pneumonia also develop bacteremia and have worse prognosis (51). Thus, we first tested host protection against invasive infection using the well-characterized serotype 4 clinical isolate *S. pneumoniae* TIGR4 as a model of pneumonia that results in bacteremia (52, 53).

At week 4 following vaccination (Fig. 1B), young (2-3 months) C57BL/6 mice were infected with 5 × 10^5^ CFU of *S. pneumoniae* TIGR4 intra-tracheally (*i*.*t*.) and assessed for clinical scores and bacteremia 24 hr post-infection and overall survival over a period of 2 weeks. We found that compared to the Sham group, vaccination of mice with LEPS significantly reduced the disease severity associated with pneumococcal disease (Fig. 6A). Importantly, this reduction in disease severity was comparable with that observed in mice vaccinated with PCV13 (Fig. 6A). When we compared bacteremia across different mice groups, both the LEPS and PCV13 vaccine significantly reduced the bacterial burden in the blood (by approximately 1,000-fold) compared to the Sham group (Fig. 6B). The reduction in blood bacterial burden seen with the LEPS formulation was also significantly lower than the Empty LEPS group (Fig. 6B). Interestingly, the PCV13 vaccinated group had significantly higher incidence of bacteremia (*p*=0.002 by Fisher’s exact test), where approximately 29% of mice became bacteremic compared to 0% of mice in the LEPS vaccinated group (Fig. 6B). When we tracked overall survival, we found that all the mice from the Sham group succumbed to pneumococcal infection within 48 hrs of infection (Fig. 6C). Mice from the Empty LEPS group showed similar kinetics with more than 50% having died within 48 hrs of challenge with *S. pneumoniae* with only 14% survival (1/7) by the end of the 2-week observation period (Fig. 6C). Vaccination with PCV13 significantly improved the survival rate of mice with only 25% of mice succumbing to infection (Fig. 6C). Strikingly, mice immunized with LEPS showed 100% survival (Fig. 6C), mirroring the effect of this vaccination on clinical score (Fig. 6A) and bacteremia (Fig. 6B). Overall, these findings highlight the high efficacy of the LEPS vaccine in preventing invasive pneumococcal infection.

**Figure 6.**
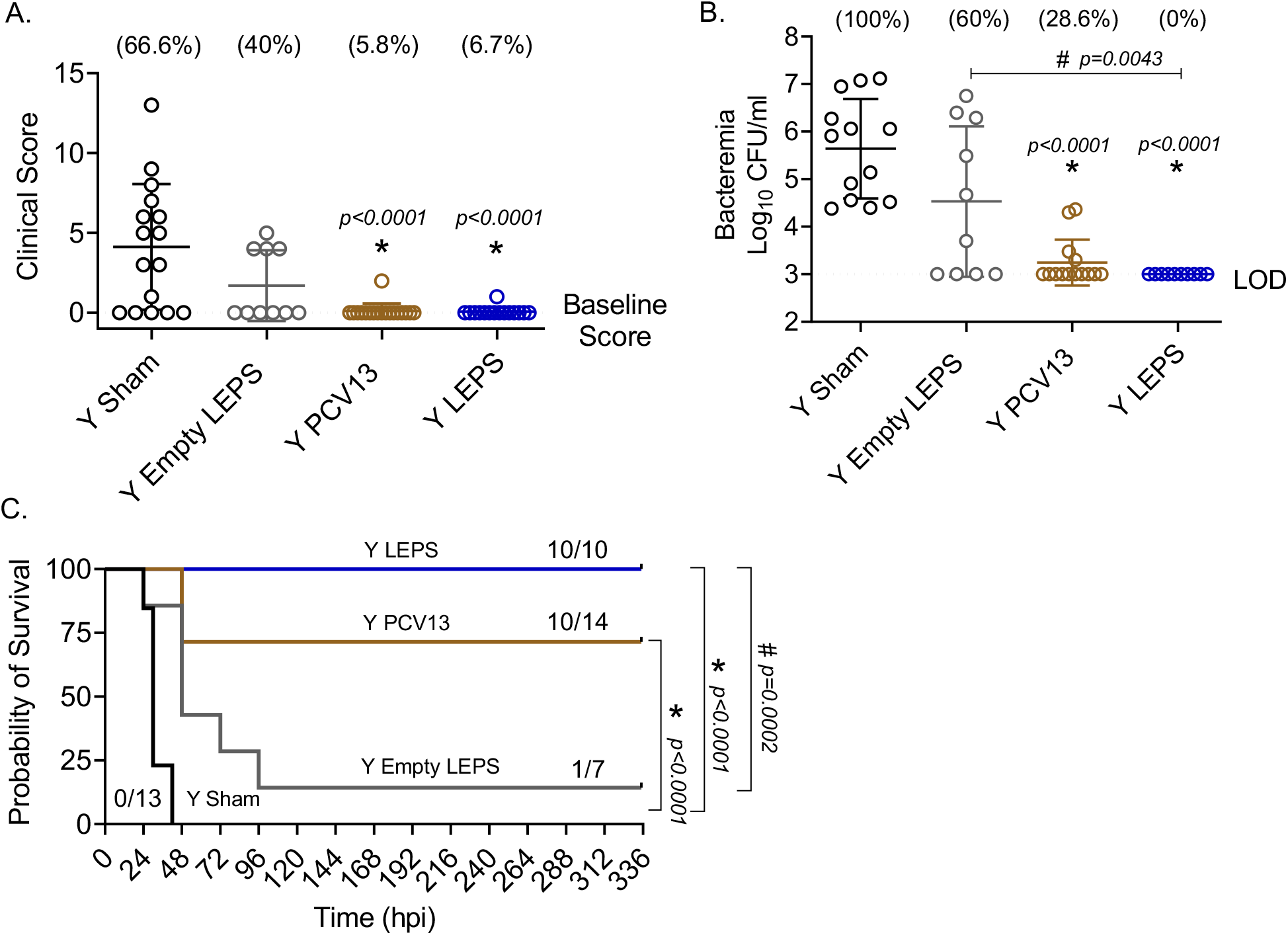
LEPS vaccination confers similar protection to Prevnar13 in young mice against invasive pneumococcal infection. Young (2-3 months) C57BL/6 mice were immunized with LEPS containing serotype 4 capsular polysaccharide antigen, PCV13, or the controls (Sham or Empty LEPS vector). Four weeks later (timeline presented in Fig. 1B), mice were infected *i*.*t* with 5 × 10^5^ CFU of *S. pneumoniae* TIGR4 strain. Clinical scores (A) and bacterial numbers in the blood along with the incidence of bacteremia (% noted above each group) (B) were determined 24 hr post-infection. Survival was also followed over time (C). (A-B) Asterisks (*) indicate significant differences with respect to the Sham group, and hash signs (#) indicate significant differences between the indicated groups as calculated by one-way ANOVA followed by Tukey’s test. Pooled data are presented as mean ± SD and each dot represents one mouse. (C) Asterisks (*) indicate significant differences with respect to the Sham group, and hash signs (#) indicate significant differences between the indicated groups as determined by the log-rank (Mantel-Cox) test. Fractions denote surviving mice. Pooled data from two separate experiments with n=13 mice in the Sham group, n=7 mice in the Empty LEPS group, n=14 mice in the PCV13 group, and n=10 mice in the LEPS group are shown. LOD: limit of detection.

### 3.6 LEPS vaccination protects old hosts against invasive pneumococcal infection

We next compared the ability of the LEPS and PCV13 vaccines in protecting aged hosts against invasive pneumococcal infection. Four weeks following vaccination (Fig. 1B), old (18-22 months) mice were infected with 5 × 10^5^ CFU of *S. pneumoniae* TIGR4 intra-tracheally (*i*.*t*.) and assessed for clinical scores and bacteremia 24 hr post-infection and survival over time. Mice belonging to both the control groups (Sham and Empty LEPS) showed severe signs of clinical disease (Fig. 7A). Although vaccination with PCV13 mitigated the overall clinical severity of disease to some extent, half of the mice from this group still experienced clinical symptoms (indicated by high clinical scores) (Fig. 7A). However, mice immunized with LEPS showed no clinical symptoms at 24 hr post-infection (Fig. 7A). When we compared bacteremia at 24 hr post-infection, we found that mice from both control groups (Sham and Empty LEPS) had a high blood bacterial burden with 100% incidence of bacteremia (Fig. 7B). Compared to the young counterparts (Fig. 6B), aged-mice experienced 100-fold higher bacterial burden in the circulation, displaying the expected age-associated susceptibility to *S. pneumoniae* infection (35, 54). Although vaccination of old mice with PCV13 significantly reduced the bacterial burden in blood compared to the Sham group, 80% of the animals still became bacteremic (Fig. 7B). Interestingly, mice immunized with LEPS had more than 1,000-fold reduction in blood bacterial numbers (Fig. 7B) compared to both control groups. Importantly, protection against bacteremia elicited by LEPS was significantly better than PCV13 as the LEPS group had a 100-fold lower bacterial burden in the blood and a lower incidence of bacteremia (*p*=0.0055 by Fisher’s exact test) where only 11% of mice became bacteremic (Fig. 7B). When we compared survival, we found that mice in both the control groups succumbed to pneumococcal infection within 24 hrs of infection (Fig. 7C). Interestingly, PCV13 failed to fully protect old mice, where we observed only 25% of animals surviving the course of infection, albeit being significantly higher than survival observed in the Sham group (Fig. 7C). In contrast, we found that the majority of mice immunized with LEPS (75%) survived the infection which was not only higher than both control groups but also significantly higher that the survival of the PCV13 group (Fig. 7C). These data indicate that while PCV13 fails to protect aged hosts, the LEPS vaccine significantly reversed the age-related susceptibility to invasive pneumococcal infection.

**Figure 7.**
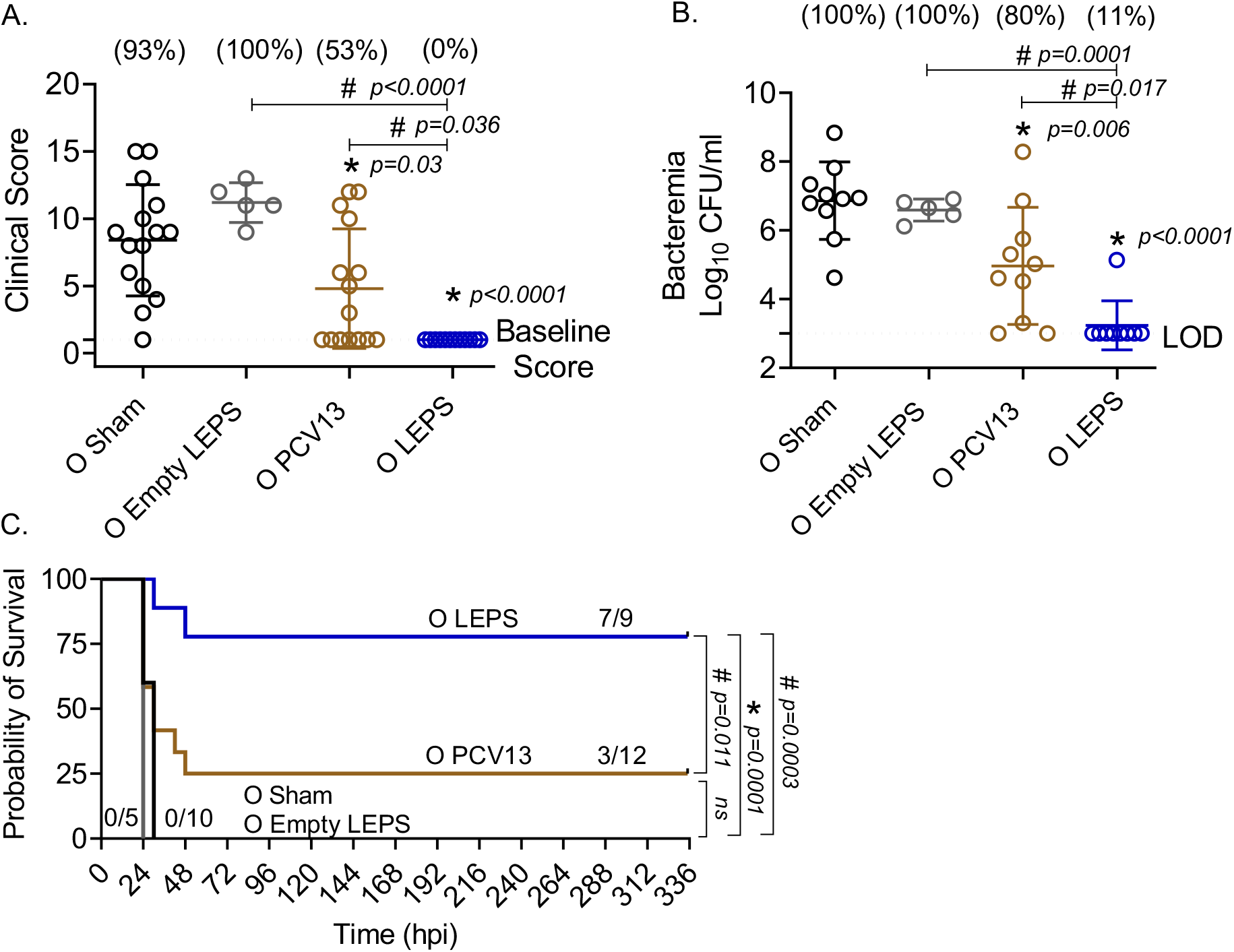
LEPS vaccination protects old mice against invasive pneumococcal infection. Old (18-22 months) C57BL/6 mice were immunized with LEPS containing serotype 4 capsular polysaccharide antigen, PCV13, or the controls (Sham or Empty LEPS vector). Four weeks later (timeline presented in Fig. 1B), mice were infected intra-tracheally with 5 × 10^5^ CFU of *S. pneumoniae* serotype 4 TIGR4 strain. Clinical scores (A) and bacterial numbers in the blood along with the incidence (% noted above each group) of bacteremia (B) were determined 24 hr post-infection. Survival was also followed over time (C). (A-B) Asterisks (*) indicate significant differences with respect to the Sham group, and hash signs (#) indicate significant differences between the indicated groups as calculated by one-way ANOVA followed by Tukey’s test. Pooled data are presented as mean ± SD and each dot represents one mouse. (C) Asterisks (*) indicate significant differences with respect to the Sham group (ns: not significant), and hash signs (#) indicate significant differences between the indicated groups as determined by the log-rank (Mantel-Cox) test. Fractions denote surviving mice. Pooled data from two separate experiments with n=10 mice in the Sham group, n=5 mice in the Empty LEPS group, n=12 mice in the PCV13 group, and n=9 mice in the LEPS group are shown. LOD: limit of detection.

### 3.7 LEPS vaccination protects old hosts against pneumococcal pneumonia

The efficacy of pneumococcal vaccines, particularly against pneumonia, decline with aging (18, 42). Therefore, we wanted to test the protective capacity of the LEPS vaccine against non-bacteremic pneumonia in aged hosts. To do so, we used a *S. pneumoniae* serotype 19F strain that is less invasive and covered by PCV13 (55). C57BL/6 old (18-22 months) mice were injected *i*.*m*. with the LEPS vaccine containing serotype 19F capsular polysaccharide antigen, PCV13, or the Sham control (Fig. 1B). At week 4 post immunization (Fig. 1B), mice were infected (*i*.*t*.) with 2 × 10^7^ CFU of *S. pneumoniae* 19F and assessed for clinical severity of disease and bacterial burden in the lung 24 hr post-infection. Interestingly, mice vaccinated with PCV13 showed worsening of clinical scores similar to that of the Sham group (Fig. 8A). In contrast, LEPS immunized mice showed an overall significantly reduced clinical score compared to both the Sham control and PCV13 groups (Fig. 8A). When we measured the lung bacterial burden, we observed a similar trend, with PCV13 failing to reduce the bacterial numbers compared to the Sham group (Fig. 8B), while mice that received the LEPS vaccine had approximately 10-fold less bacterial burden in the lungs compared to the Sham controls (Fig. 8B). Overall, these findings indicate that while vaccination with PCV13 failed to provide any protection in old mice against pneumococcal pneumonia, the LEPS vaccine boosted the ability of aged hosts to control lung infection.

**Figure 8.**
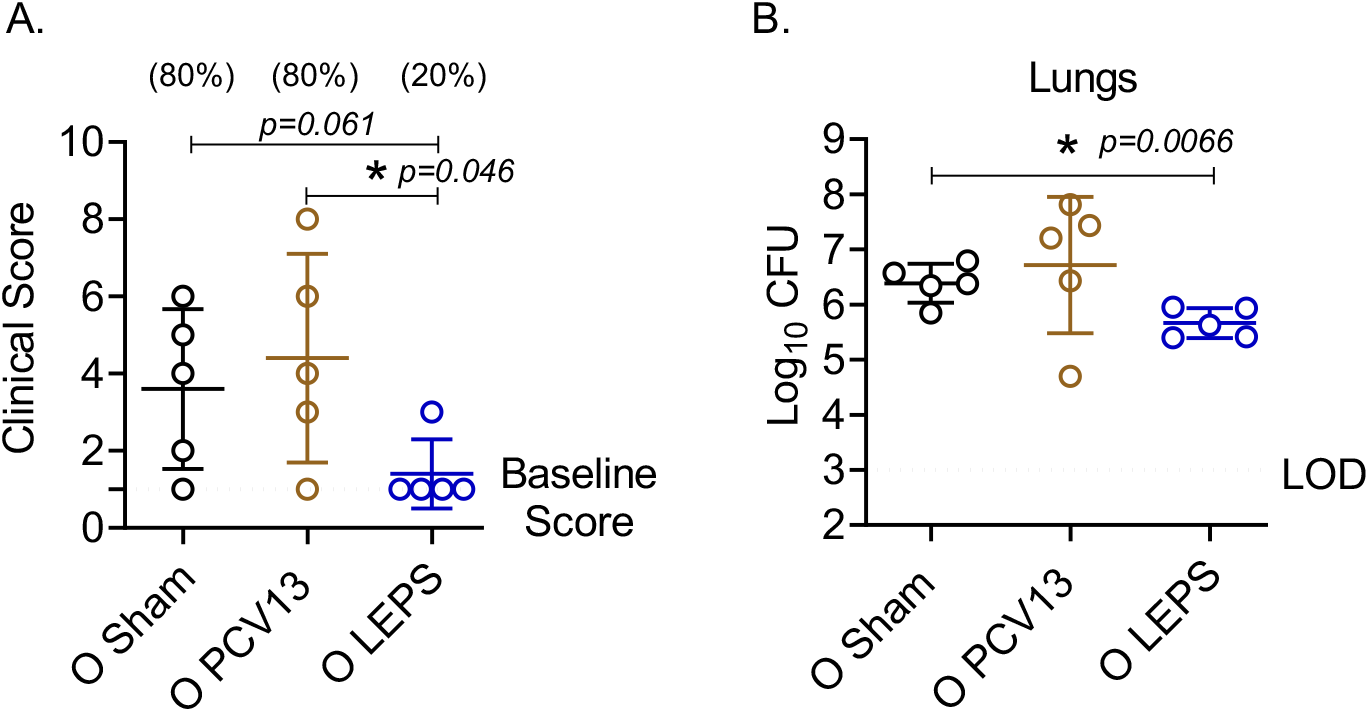
LEPS vaccination protects old mice against pneumococcal pneumonia. Old (18-22 months) C57BL/6 mice were immunized with LEPS containing serotype 19F capsular polysaccharide antigen, PCV13, or the controls (Sham or Empty LEPS vector). Following the timeline presented in Fig. 1B, 4 weeks after vaccination, mice were infected *i*.*t* with 2 × 10^7^ CFU of *S. pneumoniae* serotype 19F. Clinical scores (A) and bacterial burden in the lungs (B) were determined 24 hr post-infection. (A) Percentage of mice that displayed clinical signs of pneumonia are in parentheses. (A-B) Asterisks (*) indicate significant differences between the indicated groups as determined by one-way ANOVA followed by Tukey’s test. Data shown are pooled from n= 5 mice per group and presented as mean ± SD. Each dot represents one mouse. LOD: limit of detection.

## 4. Discussion

The current study began with the assessment of the LEPS platform in young mice to first establish protocols and confirm prior success before shifting to aged hosts. Our findings (Figs. 2, 4-6) support that the LEPS approach is a viable vaccine strategy against *S. pneumoniae* when compared to prior studies from our working group (24, 25). Though in this case, we note two differences to previous efforts. First, we use a different species of mice (C57BL/6 in this study vs. CD-1 mice in prior work). This is relevant as it suggests that the LEPS vaccine platform is effective across different mouse strains with varying genetic backgrounds, supporting future translational efforts across a diverse human population. Second, in the current study, we used the virulent-specific pneumococcus antigen PncO as an effective replacement for CRM197 in comparisons of the LEPS formulation to PCV13. Here, PncO appears to serve a very similar role to CRM197 (i.e., an immunogenic carrier protein) in promoting LEPS vaccine effectiveness both in terms of inducing a functional antibody response (Figs. 2, 4 and 5) as well as host protection against invasive pneumococcal disease (Fig. 6). Similar to prior studies (24, 25), we observed isotype switching to IgG upon vaccination with the modified LEPS that was comparable to Prevnar-13, indicating that PncO matches CRM197 in the capability to induce a T-cell dependent response (24). Since PncO serves another crucial purpose in the overall LEPS design (as discussed below), the refined formulation presented here which eliminates the need for CRM197 offers a more economical vaccine strategy against pneumococcal infections.

As pneumococcal disease has a disproportional impact upon the elderly, success of the LEPS platform in young mice, both within the current study and as reported previously (24, 25), thus prompted efforts with aged mice. However, a significant unknown was whether the LEPS platform would perform within aged hosts as the efficacy of vaccines is known to decline with aging (40, 41). PPSV is traditionally recommended for the elderly while PCV is recommended for the most vulnerable elderly with underlying conditions (56). Yet, aging leads to defects in T cell-dependent and -independent antibody production (57, 58), limiting the efficacy of both vaccines (56, 57, 59). This was recapitulated in this study where Prevnar-13 vaccination of old mice induced antibodies with subpar function (Figs. 4 and 5) resulting in reduced protection against infection (Figs. 6 and 7) as compared to young controls. In contrast, the LEPS vaccine was equally effective across host age (Figs. 6 and 7). LEPS immunization of old mice resulted in comparable antibody levels to Prevnar-13 (Fig. 3) but improved antibody function (Figs. 4 and 5). This translated to improved protection against pneumococcal infection (Figs. 7 and 8) relative to Prevnar-13. Importantly, LEPS immunization protected old mice (clinical score, bacteremia, and overall survival-Figs. 7 and 8) to a similar degree as it did young mice, suggesting that the LEPS platform is able to overcome the age-driven decline in immune responses. The detailed mechanisms by which LEPS (and its myriad of formulation variables) boosts antibody responses in aged hosts and whether it is directly activating B and/or T cells or acting as an adjuvant to boost antigen uptake and presentation by antigen presenting cells is an avenue for future studies.

An important finding here is that the LEPS platform not only provided protection against invasive pneumococcal disease (Fig. 7) but also against pneumococcal pneumonia (Fig. 8). This is of high clinical relevance as one of the limitations of licensed pneumococcal vaccines is their reduced efficacy against pneumonia in the elderly (60, 61). This is largely driven by the fact that current polysaccharide vaccines fail to account for bacterial disease progression. *S. pneumoniae* typically reside as a commensal biofilm within the human nasopharynx and asymptomatic colonization is thought to be a prerequisite of disease (16, 62). The transition from benign colonizer to lethal pulmonary or systemic pathogen involves changes in transcription profiles and bacterial phenotype (22), including changes to the surface polysaccharide capsule, which is the target of current vaccines. The LEPS vaccine is built specifically to address this weakness in current vaccine options that only focus on polysaccharide immunogens as the LEPS vehicle also includes PncO (25, 63, 64), an *S. pneumoniae* surface protein antigen (conserved across serotypes) over-represented on those virulent pneumococci that break free of the asymptomatic nasopharynx biofilm, disseminate to other bodily locations (lung, blood), and promote disease. We have yet to fully assess the functionality of the associated PncO protein in aged hosts. As this protein antigen becomes critical during later stages of pneumococcal disease, including those prompted by viral co-infection, we anticipate its relevance to emerge in secondary bacterial pneumoniae studies spurred by viral exposure in aged mice. We intend to pursue such studies in the future and, as warranted, further refine the protein antigen content of LEPS based on *S. pneumoniae* antigens that are specifically upregulated during infection of aged hosts.

In summary, this study establishes the use of the LEPS platform as a viable, effective, and economical vaccine strategy against pneumococcal infection in aged hosts. Many features of LEPS can be readily adjusted, including polysaccharide coverage and content level, the noncovalent attachment mechanism of the associated protein, and the base lipid composition. As such, we expect future optimization of the LEPS formulation to further build upon the results obtained in this work. The protein component of LEPS, in particular, holds extended potential as several protein antigens can be combined into one LEPS system for a multiplier effect in valency and for serotype-independent protection. As the elderly are projected to reach two billion worldwide by 2050 (12), the LEPS platform provides a timely intervention against serious pneumococcal infections that can be easily adapted to target other respiratory-tract pathogens that infect this vulnerable population.

## 6. Author contributions

MB conducted the research, analyzed data, and wrote the paper. RN conducted research and analyzed data. EYIT, DP, AA, and SRS conducted research. BAP and ENBG designed and supervised the research and wrote and edited the paper. All authors read and approved the final manuscript.

## 8. Acknowledgements

We would like to acknowledge Sydney Herring for critical reading and discussion of the manuscript.

## Supplemental Materials

**Figure S1. LEPS and Prevnar-13 vaccination induce comparable antibody production against *S. pneumoniae***. C57BL/6 young (2 months) and old (18-22 months) mice were injected *i*.*m*. with LEPS containing serotype 4 capsular polysaccharide antigen, PCV13, or the controls (Sham or Empty LEPS vector) following the timeline presented in Fig.1B. Levels of IgM and IgG in the sera against heat killed (HKB) *S. pneumoniae* TIGR4 were measured by ELISA. Antibody units were determined based on a hyperimmune standard included in each ELISA plate. Data shown are presented as the mean ± CI and are pooled from 3 separate experiments. For the young groups (A-B), data from n=10 mice for Empty LEPS, n=10 mice for LEPS, and n=15 mice for PCV13 are pooled. For the old groups (C-D), data from n= 5 mice for Empty LEPS, n=10 mice for LEPS, and n=10 mice for PCV13 are pooled.

