## Supplementary figures and images for "Liposomal encapsulation of polysaccharides (LEPS) as an effective vaccine strategy to protect aged hosts against *S. pneumoniae* infection"

### Supplemental Figure 1

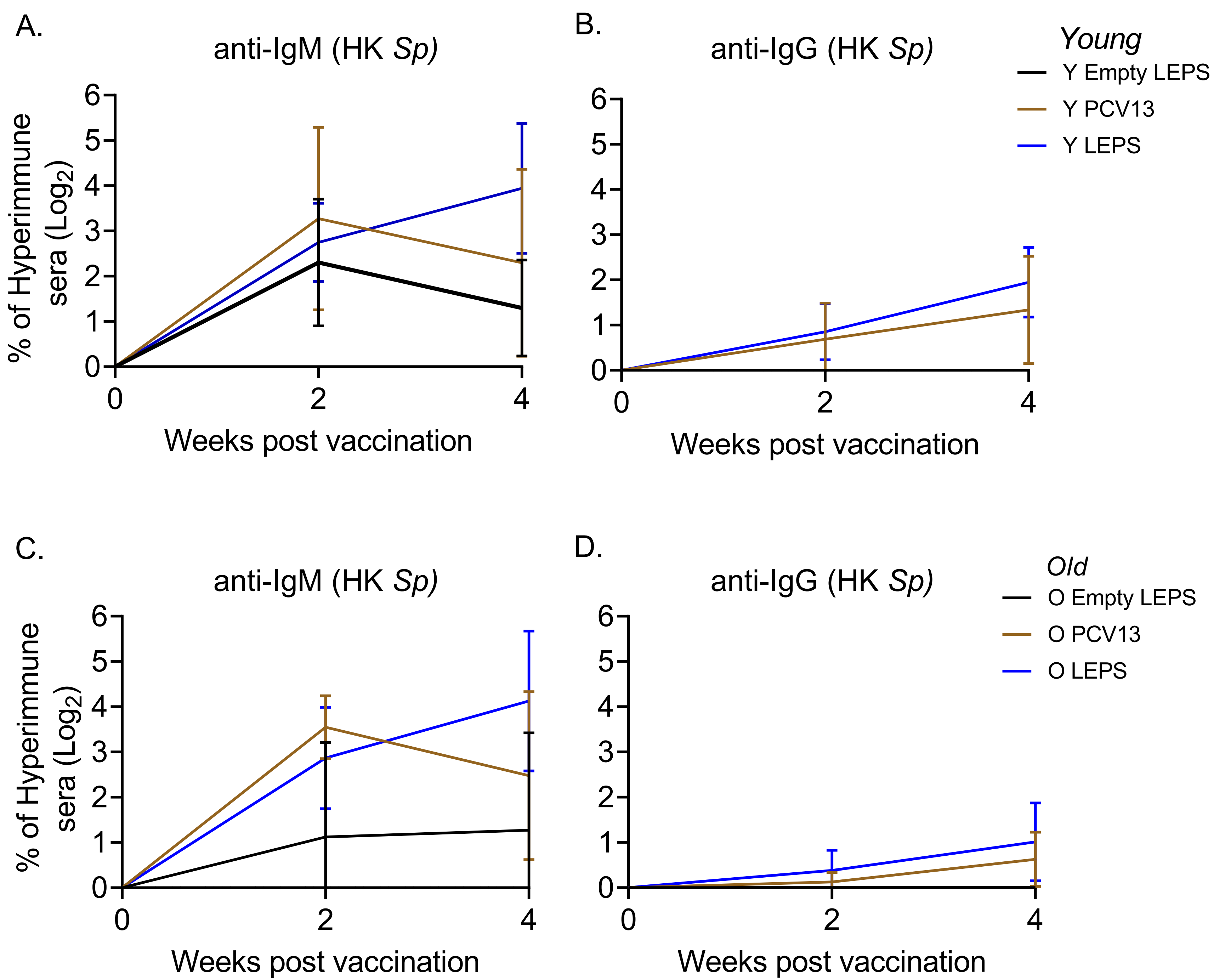
